# Comparison of three variant callers for human whole genome sequencing

**DOI:** 10.1101/461798

**Authors:** Anna Supernat, Oskar Valdimar Vidarsson, Vidar M. Steen, Tomasz Stokowy

## Abstract

Testing of patients with genetics-related disorders is in progress of shifting from single gene assays to gene panel sequencing, whole-exome sequencing (WES) and whole-genome sequencing (WGS). Since WGS is unquestionably becoming a new foundation for molecular analyses, we decided to compare three currently used tools for variant calling of human whole genome sequencing data. We tested DeepVariant, a new TensorFlow machine learning-based variant caller, and compared this tool to GATK 4.0 and SpeedSeq, using 30×, 15× and 10× WGS data of the well-known NA12878 DNA reference sample.

According to our comparison, the performance on SNV calling was almost similar in 30× data, with all three variant callers reaching F-Scores (i.e. harmonic mean of recall and precision) equal to 0.98. In contrast, DeepVariant was more precise in indel calling than GATK and SpeedSeq, as demonstrated by F-Scores of 0.94, 0.90 and 0.84, respectively.

We conclude that the DeepVariant tool has great potential and usefulness for analysis of WGS data in medical genetics.

## INTRODUCTION

Next-generation sequencing (NGS) has revolutionized the way genetic laboratories and research groups operate and perform their genomic analyses. First, genetic testing of patients for hereditary disorders has shifted from single gene assays to gene panel sequencing, and then to whole-exome sequencing (WES) and whole-genome sequencing (WGS)^1–3^. Human WGS allows detection of disease causing variants in both protein encoding- and non-coding regions of the genome^4^, with the prospect of being gradually implemented as a major tool in precision medicine^5^.

An overview of the literature (Supplementary Information 1 and 2) highlights the most common applications of WGS in a medical setting (Figure 1). WGS is nowadays used for a spectrum of genetics-related disorders: in particular monogenic disorders and genomic syndromes^3^ but also a wide range of diseases with complex inheritance, such as sporadic cancer^6,7^, heart diseases^8^, respiratory tract diseases^9^, diabetes^10^ and psychiatric conditions^11^. The number of original research articles in PubMed relevant for “human whole genome sequencing” constantly rises and nearly tripled in the last 5 years (Supplementary Information 3).

**Figure 1.**
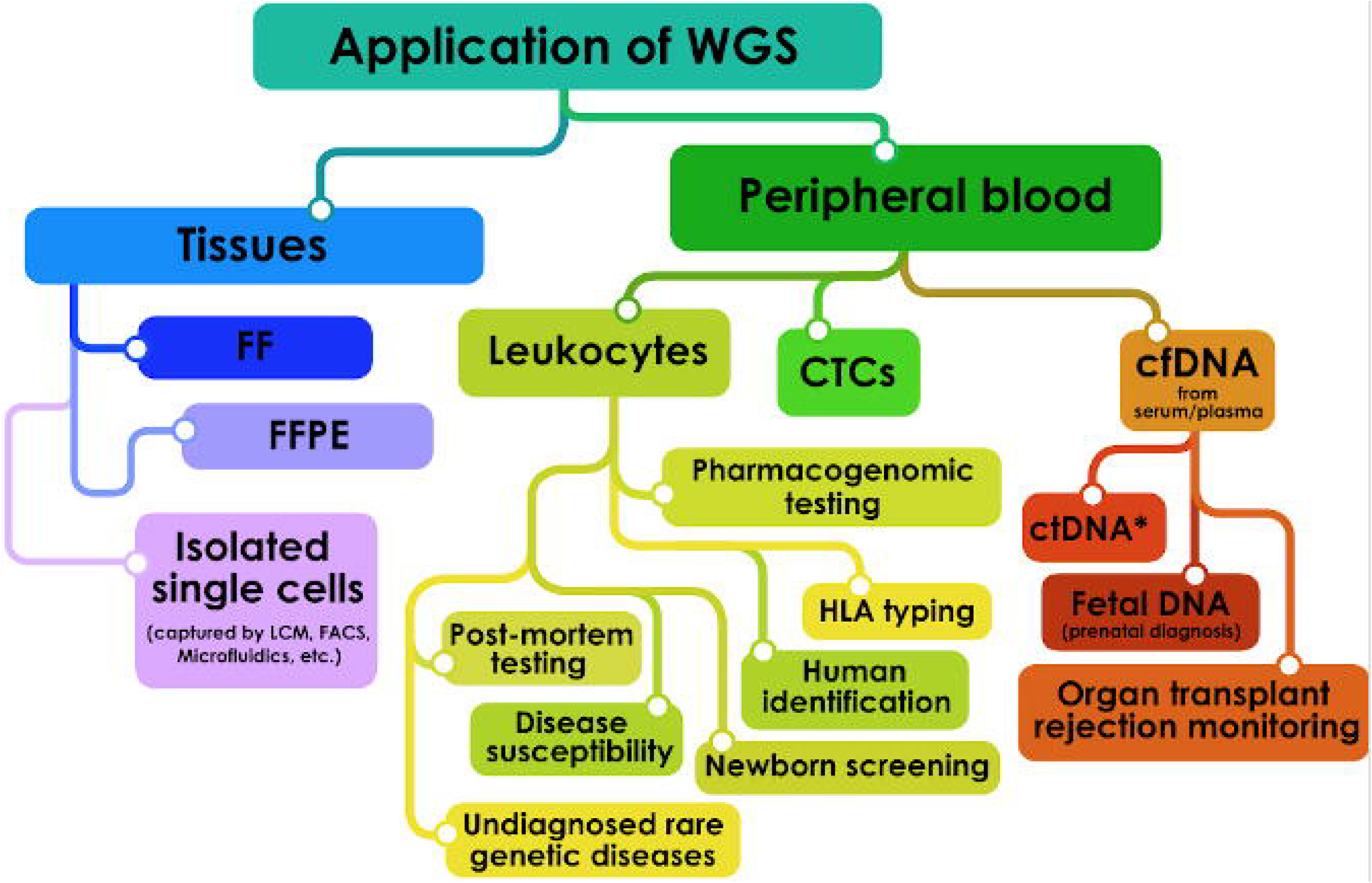
Possible applications of human whole genome sequencing (WGS) with respect to the source of biological material. Abbreviations: FF – Fresh Frozen Tissue; FFPE – Formalin Fixed Paraffin Embedded; LCM – Laser Capture Microdissection; FACS – Fluorescence Activated Cell Sorting; HLA – Human Leukocyte Antigen; CTCs – Circulating Tumor Cells; cfDNA – Circulating Free DNA; ctDNA – Circulating Tumor DNA (* detectable also in other body fluids).

However, before human WGS can become fully integrated in routine clinical diagnostics, there is an urgent need to improve and standardize the bioinformatics methods that are used in the analysis of WGS data. In general, the current workflow includes the following steps: quality control, alignment of raw data to a reference genome, variant calling (germline and/or somatic), annotation of variants, filtering of variants, data visualization and reporting (Figure 2). With respect to the types of genetic variation, single nucleotide variants (SNVs) and short indels are commonly called, whereas structural variants (SVs) and copy number variants (CNVs) have proven more challenging to detect in WGS data^12^.

**Figure 2.**
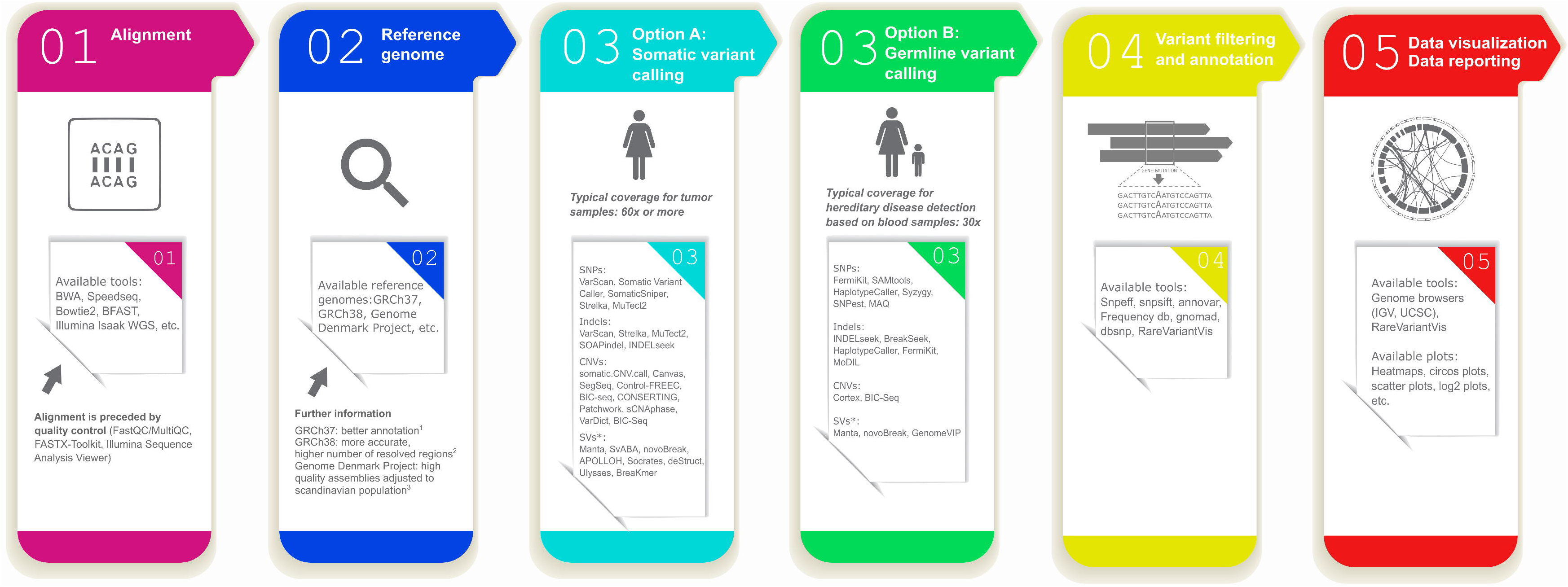
Current gold standard workflow for analysis of whole genome sequencing data.

Most studies that apply WGS data to search for genetic causes of monogenic disorders conduct variant calling by the gold standard GATK pipeline^13,14^, supported by somatic variant callers in cancer studies^15^ (see Figure 2). In this work we focus on single nucleotide variants, with the intention to evaluate structural and copy number variants in the future. Variant calling must be precise, adequate to WGS coverage and to the type of experiment. Despite recent advances in computational analysis, some parts of the workflow still require refinement. Among possible approaches towards improvement, utilization of deep learning seems to be very promising.

The most accurate variant calls for 30× human WGS data was recently reported by the PrecisionFDA Truth Challenge (https://precision.fda.gov/challenges/truth/results). The DeepVariant tool^16^ won the challenge, obtaining F-score values (i.e. harmonic mean of recall and precision) that reached 99.96% for single nucleotide variants (SNV) and 99.40% for short indels. This tool developed by the Google Brain team is the first variant calling method that applies the TensorFlow deep learning library^17^ to call variants in human genome sequencing data. To further explore the performance of this new tool, we decided to compare DeepVariant to two commonly used variant callers, namely the GATK 4.0 (the current gold standard pipeline)^13^ and SpeedSeq^18^ (a time efficient pipeline).

## RESULTS

### The performance of the DeepVariant tool in variant calling of 30× WGS data from the NA12878 DNA reference sample

In order to further explore the findings of the PrecisionFDA Truth Challenge in a real-life setting, we decided to test the performance of DeepVariant on the well-known NA12878 reference sample (sequenced in our laboratory). The sequencing resulted in 764,040,251 reads that were aligned to the GRCh38.p10 reference (99.06% of reads were aligned). The mean coverage was 34.15×, with 40.25% GC and 28.73 mean mapping quality. General sequencing error rate was 0.7% (733,229,674 base mismatches, 11,924,682 insertions and 11,666,609 deletions). The DeepVariant tool called and marked Passed Filter for a total of 4,544,442 variants, including 3,753,358 SNVs and indels: 375,878 short insertions, 399,843 short deletions (pure addition or removal of bases, according to RTG Tools manual) and 15,363 complex indels (for example length change between the reference and alternative alleles, but not pure), with transition to transversion ratio equal to 2.01 (Table 1).

**Table 1.**
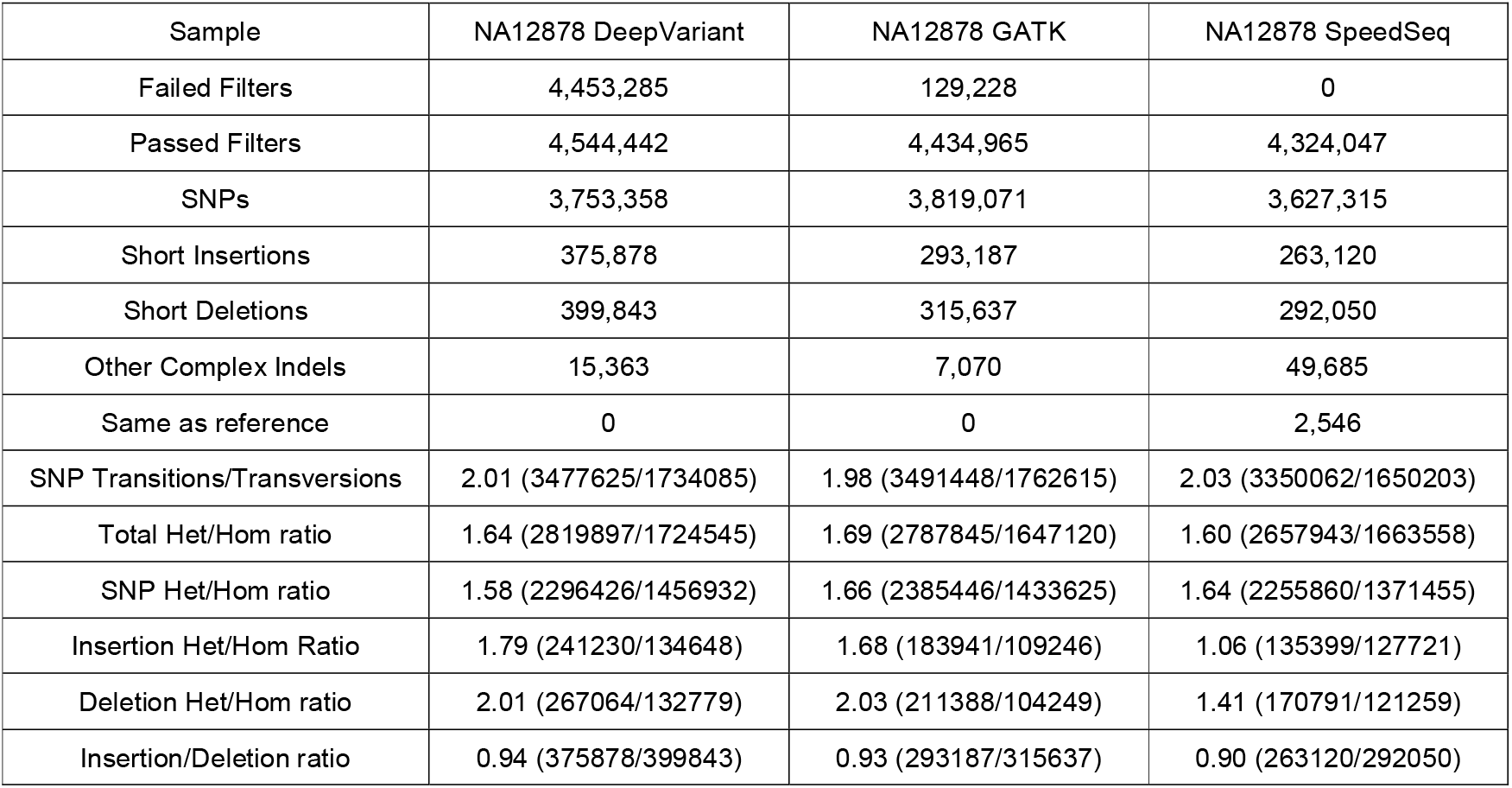
Variant calling statistics computed using RTG Tools for the three different variant calling methods. Values were computed for the raw vcf files produced by the callers.

| Sample | NA12878 DeepVariant | NA12878 GATK | NA12878 SpeedSeq |
| --- | --- | --- | --- |
| Failed Filters | 4,453,285 | 129,228 | 0 |
| Passed Filters | 4,544,442 | 4,434,965 | 4,324,047 |
| SNPs | 3,753,358 | 3,819,071 | 3,627,315 |
| Short Insertions | 375,878 | 293,187 | 263,120 |
| Short Deletions | 399,843 | 315,637 | 292,050 |
| Other Complex Indels | 15,363 | 7,070 | 49,685 |
| Same as reference | 0 | 0 | 2,546 |
| SNP Transitions/Transversions | 2.01 (3477625/1734085) | 1.98 (3491448/1762615) | 2.03 (3350062/1650203) |
| Total Het/Hom ratio | 1.64 (2819897/1724545) | 1.69 (2787845/1647120) | 1.60 (2657943/1663558) |
| SNP Het/Hom ratio | 1.58 (2296426/1456932) | 1.66 (2385446/1433625) | 1.64 (2255860/1371455) |
| Insertion Het/Hom Ratio | 1.79 (241230/134648) | 1.68 (183941/109246) | 1.06 (135399/127721) |
| Deletion Het/Hom ratio | 2.01 (267064/132779) | 2.03 (211388/104249) | 1.41 (170791/121259) |
| Insertion/Deletion ratio | 0.94 (375878/399843) | 0.93 (293187/315637) | 0.90 (263120/292050) |

### Analysis of coding sequences of the genome

Variants located within coding regions of the genome (called by the DeepVariant tool and filtered by positions of GRCh38.p10 to only include coding exons) were extracted for further evaluation. In summary, 100,687 coding variants were marked as Passed Filter, out of which 100,340 belonged to chromosomes and 347 to alternative GRCh38 contigs. Total count of coding variants included 86,145 SNVs, 7,092 short insertions, 7,256 short deletions and 194 other short indels, with transition to transversion ratio equal to 2.33.

### Comparison of the DeepVariant, GATK and SpeedSeq tools for analysis of human WGS data

DeepVariant, GATK 4.0 and SpeedSeq calls were compared to the set of NA12878 Genome in a Bottle high confidence GRCh38 variants (hosted by the National Institute of Standards and Technology, USA; NISTv3.3.2). NIST reference variants are the most reliable NA12878 variant calls available for analytical validation, thus we decided to use them in our evaluation.

Our analysis showed that DeepVariant called the highest total number of variants (4,544,442) compared to the two other interrogated tools (4,434,965 called by GATK and 4,324,047 by SpeedSeq). Still, the F-Score (i.e. harmonic mean of recall and precision, 30×) for SNVs was almost the same for DeepVariant (0.981) as compared to GATK (0.978) and SpeedSeq (0.977) (Table 2). On the other hand, DeepVariant was clearly more precise (F-Score of 0.94) in indel calling as compared to GATK and SpeedSeq (F-Scores 0.90 and 0.84, respectively). These quality scores are backed up by the highest number of true positive indel calls (460,271) as well as the lowest number of false negative (39,426) and false positive indel calls (16,122), for DeepVariant, as presented in Table 2.

**Table 2.** Comparison of variant calling pipelines. Variants were called from 30×, 15× and 10× coverage of the NA12878 sample (HiSeq 4000, Genomics Core Facility, Bergen, Norway) and compared to GIAB NISTv3.3.2 (ftp://ftp-trace.ncbi.nlm.nih.gov/giab/ftp/release/NA12878_HG001/NISTv3.3.2/GRCh38/). The GIAB true variant set included 3,042,789 SNV variants and 499,697 indels. Variant counts and performance scores were estimated using hap.py – an Illumina haplotype comparison/benchmarking tool.

| SNV | True positive SNV calls | False negative SNV calls | False positive SNV calls | Genotype mismatch | Total number of SNV calls | SNV calling precision | SNV recall | F1 Score |
| --- | --- | --- | --- | --- | --- | --- | --- | --- |
| SpeedSeq 30× | 2,942,217 | 100,572 | 38,107 | 11,869 | 3,802,913 | 0.987223 | 0.966947 | 0.97698 |
| SpeedSeq 15× | 2,814,843 | 227,946 | 57,654 | 31,131 | 3,613,466 | 0.97994 | 0.925086 | 0.951724 |
| SpeedSeq 10× | 2,589,184 | 453,605 | 84,123 | 53,955 | 3,334,440 | 0.968548 | 0.850925 | 0.905934 |
| DeepVariant 0.4.1 30× | 2,948,290 | 94,499 | 22,902 | 19,595 | 3,714,945 | 0.992294 | 0.968943 | 0.98048 |
| DeepVariant 0.4.1 15× | 2,903,519 | 139,270 | 55,261 | 41,999 | 3,674,970 | 0.981328 | 0.954229 | 0.967589 |
| DeepVariant 0.4.1 10× | 2,809,014 | 233,775 | 84,054 | 61,314 | 3,573,547 | 0.970952 | 0.923171 | 0.946459 |
| GATK 4.0 – WDL 30× | 2,952,605 | 90,184 | 41,684 | 12,579 | 3,814,443 | 0.986082 | 0.970361 | 0.978159 |
| GATK 4.0 – WDL 15× | 2,891,815 | 150,974 | 59,476 | 31,151 | 3,698,103 | 0.979851 | 0.950383 | 0.964892 |
| GATK 4.0 – WDL 10× | 2,763,913 | 278,876 | 82,452 | 57,639 | 3,526,795 | 0.971036 | 0.908349 | 0.938647 |
| INDEL | True positive INDEL calls | False negative INDEL calls | False positive INDEL calls | Genotype mismatch | Total number of INDEL calls | INDEL calling precision | INDEL recall | F1 Score |
| SpeedSeq 30× | 383,930 | 115,767 | 32,263 | 13,310 | 619,159 | 0.923499 | 0.768326 | 0.838796 |
| SpeedSeq 15× | 337,815 | 161,882 | 34,635 | 16,172 | 542,025 | 0.907915 | 0.67604 | 0.775005 |
| SpeedSeq 10× | 290,678 | 209,019 | 35,029 | 18,179 | 466,079 | 0.893253 | 0.581709 | 0.704578 |
| DeepVariant 0.4.1 30× | 460,271 | 39,426 | 16,122 | 8,147 | 816,456 | 0.967406 | 0.9211 | 0.943685 |
| DeepVariant 0.4.1 15× | 428,557 | 71,140 | 29,651 | 15,010 | 748,972 | 0.937303 | 0.857634 | 0.8957 |
| DeepVariant 0.4.1 10× | 387,075 | 112,622 | 38,695 | 20,121 | 668,593 | 0.911332 | 0.774619 | 0.837433 |
| GATK 4.0 – WDL 30× | 429,859 | 69,838 | 24,191 | 9,251 | 764,422 | 0.948269 | 0.860239 | 0.902112 |
| GATK 4.0 – WDL 15× | 380,932 | 118,765 | 30,603 | 11,918 | 655,658 | 0.927084 | 0.762326 | 0.836671 |
| GATK 4.0 – WDL 10× | 335,446 | 164,251 | 34,626 | 14,030 | 569,141 | 0.90753 | 0.671299 | 0.771742 |

With respect to the performance on WGS data with lower coverage (i.e. 15× and 10×), we observed that reduced coverage resulted in a marked drop of the quality of variant calling for all tools (Table 2). Independently of the coverage, DeepVariant was the most precise caller in all our comparisons.

Indeed, the F-Scores of DeepVariant for 15× data were almost similar to SpeedSeq at 30×. Detailed interrogation of false positive and false negative variants indicated that out of the three tested variant callers, GATK was most prone to errors in low coverage regions, while DeepVariant was most robust in such regions (Supplementary Information 4).

According to our findings, base change and context of false positive variants seemed to depend on the caller, while false negative variants appeared in the regions of lower coverage. GATK calls more A>T, C>A, G>T and T>A substitutions, than expected from the distribution of such variants in the human genome (Supplementary Information 5). SpeedSeq calls more A>C, A>T, C>A, G>T, T>A and T>G substitutions, while false positive and false negative calls by DeepVariant seem to be independent with respect to the base change.

## DISCUSSION

In this study, we confirm the results of PrecisionFDA Truth Challenge, demonstrating that the new DeepVariant tool is currently the most accurate variant caller available and therefore has great potential for implementation in routine genome diagnostics. Interestingly, this TensorFlow machine learning-based method outperforms the latest version of GATK – a gold standard method that was first published in 2010^13^. The DeepVariant algorithm takes pictures of aligned reads and then uses machine learning to decide about the presence and the type of each variant. This novel method is an interesting alternative to previously used approaches, which are mainly based on counting reads with alternative sequence in a certain genomic position (GATK, SpeedSeq and others).

The DeepVariant SNV and indel calling F1 performance scores obtained in our analysis are lower than those obtained in the FDA Challenge: 0.981 versus 0.999 and 0.94 versus 0.99, respectively. Raw data filtering and optimization of caller parameters are essential for variant calling outcome^19–21^, and to provide a reliable benchmark we decided to follow the instructions that were available on the authors websites (links are listed in the Methods section). We provide all our raw data and variant calls along with source code available for scientific community discussion.

Interestingly, DeepVariant proved to be the most precise caller, irrespectively of sequence coverage. As an example, the F-Scores obtained by DeepVariant at 15× were comparable to SpeedSeq at 30×. This suggests that the application of a high precision caller can markedly reduce the cost of sequencing consumables while keeping the same performance. Furthermore, at lower coverage, GATK and SpeedSeq would call more A>T, C>A G>T and T>A substitutions than expected from the distribution of variants in the human genome. At the same time, false positive and false negative calls by DeepVariant seemed to be independent with respect to the base change. Our statistics of such incorrectly called variants could improve the understanding the challenges of each caller and aid the development of new variant calling algorithms in future.

It is important to notice that local setup of the DeepVariant tool on an offline Unix machine was trivial when following the authors instructions: using a portable Docker container or building from source. With regards to the complexity of the computational resources for running all the tools, our experience showed that 8 core machines with 16GB RAM was the minimum hardware setting to run a WGS pipeline. In such a setting, the complete WGS analyses would usually take from 24 to 48 hours. However, it was possible to accelerate the computations: For example, the SpeedSeq pipeline on a 72 core/100GB RAM machine was run in approximately 3 hours per sample, while the DeepVariant variant calling time was reduced by more than 50% using GPU with 4 GB VRAM and CUDA support.

In summary, we conclude that TensorFlow-based variant calling in human WGS data has great potential and usefulness for medical genetics. Algorithms used by Ryan Poplin, Marc DePristo and colleagues will most likely open new, fresh perspective in genomics and bioinformatics.

## METHODS

### Whole genome sequencing, quality control and alignment of the NA12878 DNA reference sample

For the purpose of this work, we purchased the NA12878 cell line (CEPH/UTAH PEDIGREE Live Culture) from Coriell Cell Repositories (http://ccr.coriell.org/). Whole genome sequencing of this sample was performed by the Genomics Core Facility (GCF) at the University of Bergen, Norway, using an Illumina HiSeq 4000 instrument and the Illumina 150 bp TruSeq DNA PCR-FREE paired-end sequencing protocol, aiming at 30× coverage. Obtained sequences were deposited in the NCBI SRA repository under the PRJNA436473 BioSample record. We performed quality control of the raw reads with FastQC and used MultiQC to generate quality control reports for our samples. Reads were aligned to the human reference genome – Gencode GRCh38.p10^22^ using bwa-mem^23^ in a secure SAFE computational infrastructure (https://it.uib.no/SAFE). Aligned sequences were deposited in the NCBI SRA repository. The quality of the obtained bam file was evaluated using Qualimap software^24^.

### Variant calling and comparison of variant calling methods

We performed and compared variant calling using three different analysis tools: DeepVariant 0.4.1 (winner of the FDA Challenge), GATK 4.0.0.0 (the most recent standalone version of a gold standard pipeline, https://gatkforums.broadinstitute.org/wdl/categories/wdl-documentation) and SpeedSeq 0.1.0 (rapid analysis pipeline, recently developed by Chiang and colleagues^18^) to obtain SNV vcf files for our NA12878 sample. The DeepVariant analysis was performed in accordance with online instructions (https://github.com/google/deepvariant/blob/r0.5/docs/deepvariant-case-study.md). The GATK analysis was based on a best practices pipeline from The Broad Institute (https://github.com/oskarvid/wdl_germline_pipeline/tree/4.0). SpeedSeq variant calling was conducted using the SpeedSeq var command, in accordance with the instructions from the authors website (https://github.com/hall-lab/SpeedSeq). The results of all three variant calling pipelines were compared to the GIAB NISTv3.3.2 true variant set: ftp://ftp-trace.ncbi.nlm.nih.gov/giab/ftp/release/NA12878_HG001/NISTv3.3.2/GRCh38/. Obtained results were summarized in Table 2 and further evaluated using RTG-Tools (https://github.com/RealTimeGenomics/rtg-tools) and hap.py (https://github.com/Illumina/hap.py/blob/master/doc/happy.md).

### Variant filtering and annotation

Variant filtering for coding sequences of the genome was performed using bedtools intersect^25^. As a reference file for the annotation of the genomic positions of the genes, we used Gencode gtf reference version 27. Additionally, the awk unix command was applied to extract records from the gtf file which represent exons of coding genes.

## DATA ACCESS

Raw and aligned whole genome sequencing data are available in the following NCBI SRA repository: https://www.ncbi.nlm.nih.gov/Traces/study/?acc=SRP133725

Variants called using three different algorithms and a filtered list of variants are available on GitHub pages: https://github.com/tstokowy/CoriellIndex_VCF_180306

The GATK 4.0.0.0 pipeline used in this study is available on GitHub pages: https://github.com/oskarvid/wdl_germline_pipeline/tree/4.0

In this study we used the publically available GIAB NISTv3.3.2 true variant set to evaluate variant caller performance: ftp://ftp-trace.ncbi.nlm.nih.gov/giab/ftp/release/NA12878_HG001/NISTv3.3.2/GRCh38/

## ACKNOWLEDGEMENTS

We would like to acknowledge our colleagues and collaborators for help with this work: Rita Holdhus for sequencing of samples used in this study, Ove Bruland for purchase and quality control of samples from Coriell Institute, Paweł Sztromwasser for suggestions regarding the DeepVariant tool and fruitful discussions on bioinformatics related issues, Kornel Labun for contribution to the RareVariantVis R package applied in this study.

The Genomics Core Facility (GCF) at the University of Bergen, which is part of the NorSeq consortium, provided services on herewith reported Whole Genome Sequencing; GCF is supported in part by major grants from the Research Council of Norway (grant no. 245979/F50) and Bergen Research Foundation (BFS).

This work was performed in SAFE, a solution for secure processing of sensitive personal data in research managed by the IT-department at the University of Bergen. http://it.uib.no/SAFE

We would like to acknowledge IT support from Elixir Norway http://www.bioinfo.no/elixir, especially Inge Jonassen and Kjell Petersen.

## AUTHOR CONTRIBUTION

A.S. and T.S. designed and directed the project; T. S. and V.M.S. gathered data; T.S., O.V., and A.S. analysed sequencing data; A.S. drew figures and reviewed the literature, T.S. and O.V. prepared tables and performed statistical analysis, A.S., V.M.S and T.S. wrote the article, all authors read and accepted final version of the article.

